# Protein sequencing with single amino acid resolution discerns peptides that discriminate tropomyosin proteoforms

**DOI:** 10.1101/2024.11.04.621980

**Authors:** Natchanon Sittipongpittaya, Kenneth A. Skinner, Erin D. Jeffery, Emily F. Watts, Gloria M. Sheynkman

## Abstract

Protein variants of the same gene—proteoforms—can have high molecular similarity yet exhibit different biological functions. Thus, identifying unique peptides that unambiguously map to proteoforms can provide crucial biological insights. In humans, four human tropomyosin (TPM) genes produce similar proteoforms that can be challenging to distinguish with standard proteomics tools. For example, TPM1 and TPM2 share 85% sequence identity, with amino acid substitutions that play unique roles in muscle contraction and myopathies. In this study, we evaluated the ability of the recently released Platinum single-molecule protein sequencer to detect proteoform-informative peptides. Platinum employs fluorophore-labeled recognizers that reversibly bind to cognate N-terminal amino acids (NAAs), enabling polypeptide sequencing within nanoscale apertures of a semiconductor chip that can accommodate single peptide molecules. As a proof of concept, we evaluated the ability of Platinum to distinguish three main types of proteoform variation: paralog-level, transcript-level, and post-translational modification (PTM). We distinguished paralogous TPM1 and TPM2 peptides differing by a single isobaric residue (leucine/isoleucine). We also distinguished tissue-specific TPM2 spliceforms. Notably, we found that a phosphotyrosine-modified peptide displayed reduced recognizer affinity for tyrosine, showing sensitivity to PTMs. This study paves the way for the targeted detection of proteoform biomarkers at the single molecule level.

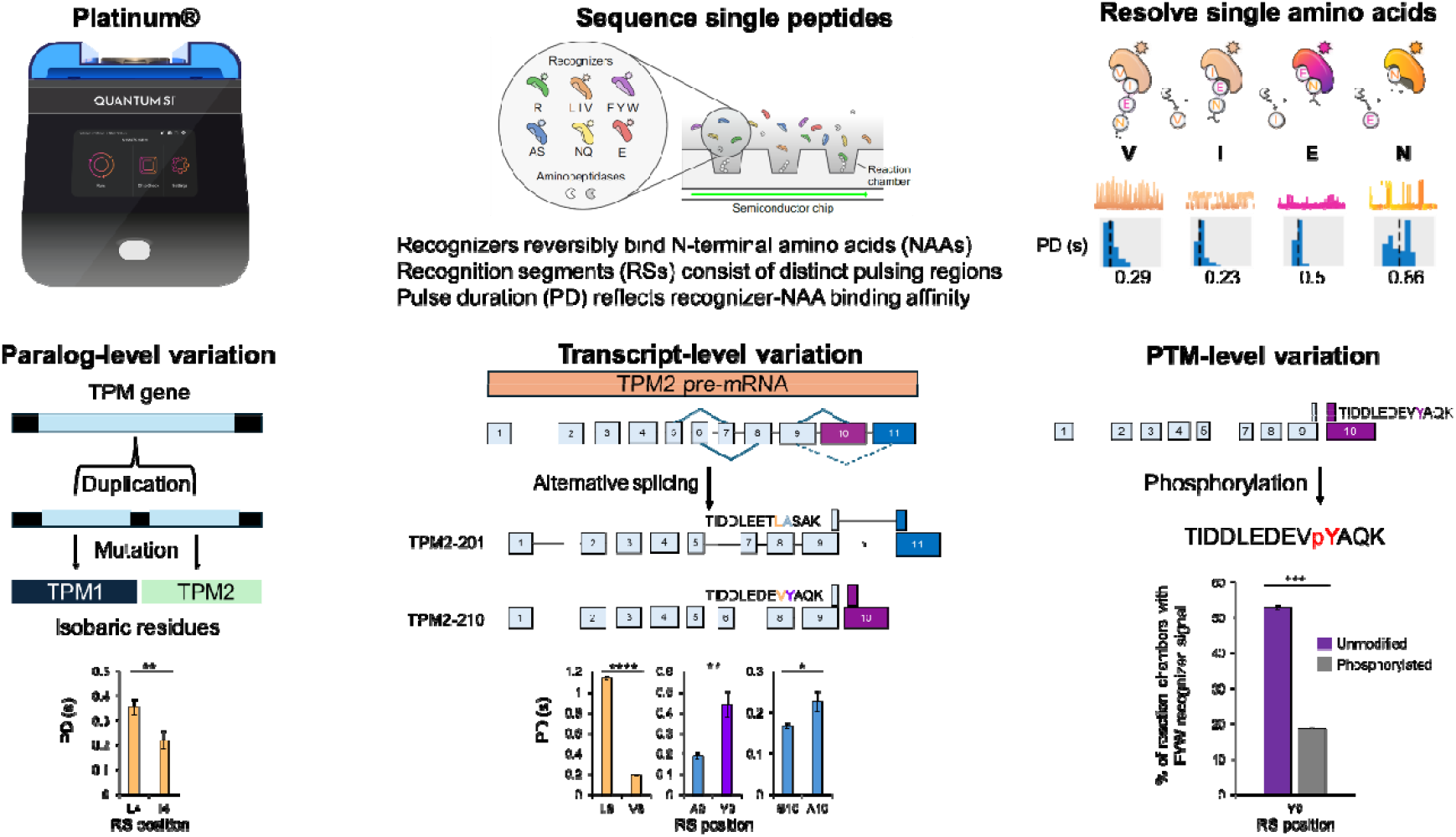

## Introduction

Proteins serve as the vital molecular bridge between genotype and phenotype, making their functional characterization critical for complete understanding of biological processes^1^. Protein function, however, is not fully encoded by the genome. Alternative splicing (AS) of pre-mRNA transcripts can produce distinct protein isoforms^2,3^. Genes can also be regulated at the post-transcriptional level, and proteins often undergo post-translational modifications (PTMs) that can alter function^4^. The protein products of genetic, post-transcriptional, and post-translational variation—collectively termed proteoforms^5^— can exhibit strikingly diverse functions that give rise to phenotypes in human health and disease^2,4^. Thus, detecting these proteoforms is key for understanding the molecular link between protein function and phenotype.

Characterizing proteoform expression can present challenges. Currently, the existence of most proteoforms is inferred from mRNA transcripts^6^ and putative PTM sites^7^. However, not all mRNA transcripts are translated into protein isoforms^8^, and some transcripts are degraded without translation^9–11^. In addition, even if putative PTM sites for all proteoforms are known, the exact proteoforms in a sample cannot be identified unless PTMs are detected and localized to specific sites^12^. Indeed, the vast majority of predicted proteoforms have yet to be validated at the peptide or protein level^13,14^. To bridge this gap, it is necessary to leverage a range of analytical technologies with varying capabilities, both independently and in combination.

Traditional antibody-based assays cannot distinguish proteoforms unless antibodies with high proteoform specificity are available^15,16^. While mass spectrometry (MS)-based methods have benefits such as high throughput and versatility, they can miss proteoform-informative peptides due data-dependent acquisition (DDA) selection bias towards the most intense peptide ions^17^. In addition, resolving peptides with similar physicochemical properties is challenging with size- and charge-based chromatographic methods^18^. Recently, there has been considerable progress in identifying peptides that uniquely indicate specific proteoforms^19,20^, an important advancement for discerning proteoforms with high amino acid sequence identity. However, to fully characterize proteoforms, it will be necessary to employ new and orthogonal detection methods capable of addressing some of the drawbacks of current methods.

Recently, several single molecule protein sequencing approaches have emerged with the potential to detect subtle amino acids and variations^21–23^. Recognizer-based sequencing, also referred to as next-generation protein sequencing (NGPS), was the first of these to be commercialized with the release of the Platinum instrument (Quantum-Si). Therefore, we sought to evaluate the feasibility of distinguishing proteoform-informative peptides using NGPS on Platinum. For this study, we chose a model proteoform family—tropomyosins (TPMs)—that shares high amino acid sequence similarity (**Figure 1**) and represents diversity at the levels of genetic polymorphisms, RNA splice variants, and PTMs^30^.

**Figure 1.**
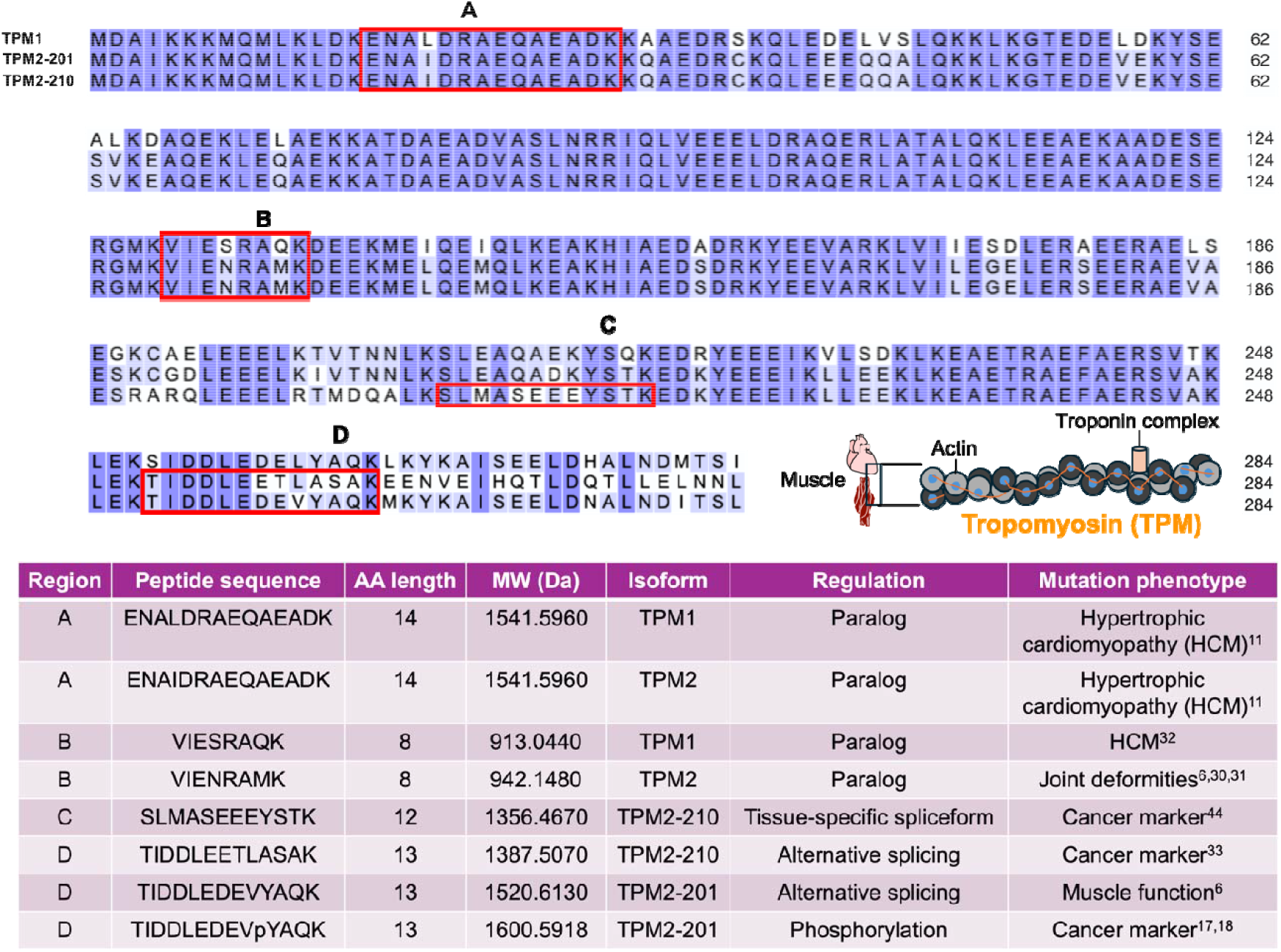
Sequence alignment of TPM1 and TPM2 isoforms and relevance of amino acid mutations to pathology. The primary structures of canonical TPM1 and two TPM2 spliceforms are shown. TPM and troponin associate with actin to form the contractile apparatus. Region A corresponds to a N-terminal region that contains isobaric peptides that differ by leucine (Leu; L) or isoleucine (Ile; I). Mutations in these paralogs have been linked with hypertrophic cardiomyopathy (HCM)^24^. Region B delineates an internal eight amino acid long region within TPM1/2 paralogs. Mutations within this region are also found in HCM and joint deformities due to Sheldon-Hall syndrome and distal arthrogryposis^25–27^. Region C highlights an isoform-informative peptide (SLMASEEEYSTK) generated via alternative splicing. Region D maps to an alternatively spliced C-terminal region of TPM2-201 and TPM2-210. While TIDDLEETLASAK (TPM2-201) and TIDDLEDEVYAQK (TPM2-210) share the same amino acid length, TIDDLEDEVYAQK is subject to phosphorylation on a single tyrosine residue^28,29^.

Filament proteins such as TPMs are among the most highly conserved proteins in evolution, playing critical roles in cellular architecture, cytoskeletal movement, and compartmental trafficking^30^. TPMs are alpha-helical coiled coil dimers that regulate the stability of actin filaments in muscle and non-muscle cells^31^. Via alternative splicing, tissue-specific promoter usage, and different poly (A) addition sites, the four human TPM genes (**Supplementary Figure 1**) collectively produce more than 40 mRNA variants that can be translated as tissue-specific proteins in skeletal and non-skeletal tissue types^30,32^.

Missense mutations in TPMs are linked to cardiac diseases such as hypertrophic cardiomyopathy (HCM)^24^ and skeletal muscle diseases^24,33–35^ (**Figure 1**). These mutations may modulate interactions of TPMs with other filament proteins, impacting the contractile apparatus^36–38^. Altered TPM expression and phosphorylation have also been linked with dilated cardiomyopathy (DCM) and cancer phenotypes^28,29^. Because TPMs are subject to extensive variation at the genetic, post-transcriptional, and post-translational levels^39^, the functional diversity and high sequence similarity of TPM proteoforms presents a daunting challenge with current analytical methods^40^.

Therefore, in this study, we sought to determine whether Platinum could distinguish between diverse TPM proteoforms. We focused on three major types of variants: isobaric peptides from TPM1/2 gene paralogs, peptides derived from tissue-specific TPM2 spliceforms, and phosphotyrosine-modified variants from unmodified peptides. In the process, we leveraged key features of the Platinum sequencing data type: characteristic pulse durations (PDs), which distinguish structurally similar NAAs, and the order of discrete recognition segments (RSs), which pinpoint amino acid location. Together, these measurements produce “kinetic signatures” that provide information on both amino acid identity and sequential order. Given that sequence variations within our model system correspond to regions relevant to TPM pathophysiology (**Figure 1**), we envision broader applications of NGPS to the targeted detection of proteoform-informative peptides from biological samples.

## Materials and Methods

### In silico digestion and peptide selection criteria

Protein sequences of canonical TPM1 and TPM2 spliceforms^41^ (**Figure 1**) were extracted from GENCODE (v43). In-silico LysC^42^ digestion of TPM sequences was performed using a custom R script leveraging the R package cleaver. The resulting peptide sequences were mapped to their corresponding genomic coordinates using PoGo and visualized as a UCSC browser track to identify peptides that are specific to TPM1 (ENALDRAEQAEADK and VIESRAQK) or TPM2 (ENAIDRAEQAEADK and VIENRAMK), shared between spliceforms of TPM2 (VIENRAMK), and specific to certain spliceforms of TPM2 (SLMASEEEYSTK, TIDDLEETLASAK, and TIDDLEDEVYAQK).

Peptides that are paralog-specific, shared between TPM2 spliceforms, and TPM2 spliceform-specific were filtered according to the following criteria: the peptide contains a C-terminal lysine (Lys; K), the peptide contains ≥ 3 amino acids that are cognate to the NAA recognizers, the peptide sequence can be recognized by ≥ 3 different recognizers (**Supplementary Figure 3**), and the peptide contains 5-25 amino acids. These criteria are important for successful sequencing of peptides by NGPS (Library Preparation Kit -Lys-C Data Sheet and Platinum Analysis Software Data Sheet).

### Peptide synthesis

Eight peptides identified from the *in silico* peptide screening (**Figure 1**) were synthesized by JPT Peptide Technologies, Inc. (Berlin, Germany) with C-terminal carboxylic acid and azido-lysine modifications (1 mg, >90% purity, lyophilized). Peptides were reconstituted in 50% acetonitrile to a concentration of 5 mM each and stored at -80°C.

Synthetic phosphopeptides and unphosphorylated peptides with C-terminal azido-lysine modifications were synthesized by Genscript (Piscataway, NJ, USA) (4 mg, >90% purity, lyophilized). The peptides were reconstituted in dimethylformamide to a concentration of 5 mM each and stored at -80°C.

### LC-MS/MS analysis

Peptide separation was performed using nanoflow high-performance liquid chromatography (HPLC) on a Dionex Ultimate 3000 system (Thermo Fisher Scientific, Bremen, Germany). Peptides were initially loaded onto an Acclaim PepMap 100 trap column (300 μm × 5 mm, 5 μm C18), followed by gradient elution through an Acclaim PepMap 100 analytical column (75 μm × 25 cm, 3 μm C18) for enhanced separation. Mass spectrometry (MS) analysis was conducted using an Orbitrap Eclipse Tribrid mass spectrometer (Thermo Fisher Scientific, Bremen, Germany) equipped with the Orbitrap Eclipse Tune (version 4.0.4091) and Xcalibur software (version 4.5.445.18) for data acquisition and analysis.

### Preparation of linker-conjugated peptide libraries

Synthetic peptides with C-terminal azido-lysine modifications were conjugated to linker molecules via strain-induced click conjugation^43^. Peptides were diluted to a final concentration of 50 µM in 100 mM HEPES, pH 8.0 (20% acetonitrile) and incubated overnight with 2 µM linker at 37 °C. The resulting conjugated peptide libraries were stored at -20 °C until sequencing.

### Peptide sequencing on Platinum

Experiments were conducted in accordance with Library Preparation Kit -Lys-C Data Sheet and Platinum Instrument and Sequencing Kit V2 Data Sheet (February 27, 2024).

Library Preparation Kit -Lys-C Data Sheet link: https://www.quantum-si.com/resources/product-data-sheets/library-preparation-kit-lys-c-data-sheet/

Platinum Instrument and Sequencing Kit V2 Data Sheet link: https://www.quantum-si.com/resources/product-data-sheets/platinum-instrument-and-sequencing-kit-v3-data-sheet/

Briefly, conjugated peptides are immobilized in nanoscale reaction chambers on a semiconductor chip (**Figure 2A**) for exposure to a mixture of freely diffusing NAA recognizers and aminopeptidases. The mixture consisted of NAA recognizers that target 12 of 20 canonical AAs (**Figure 2B**). During on-chip sequencing, NAA recognizers reversibly bind cognate NAAs, producing recognition segments (RSs) that are captured by the semiconductor chip (**Figure 2C**). To carry out the sequencing process, aminopeptidases cleave the peptide bond and expose the subsequent NAA for recognition. After 10 hours of runtime, sequencing data is transferred to the Platinum Analysis Software.

**Figure 2.**
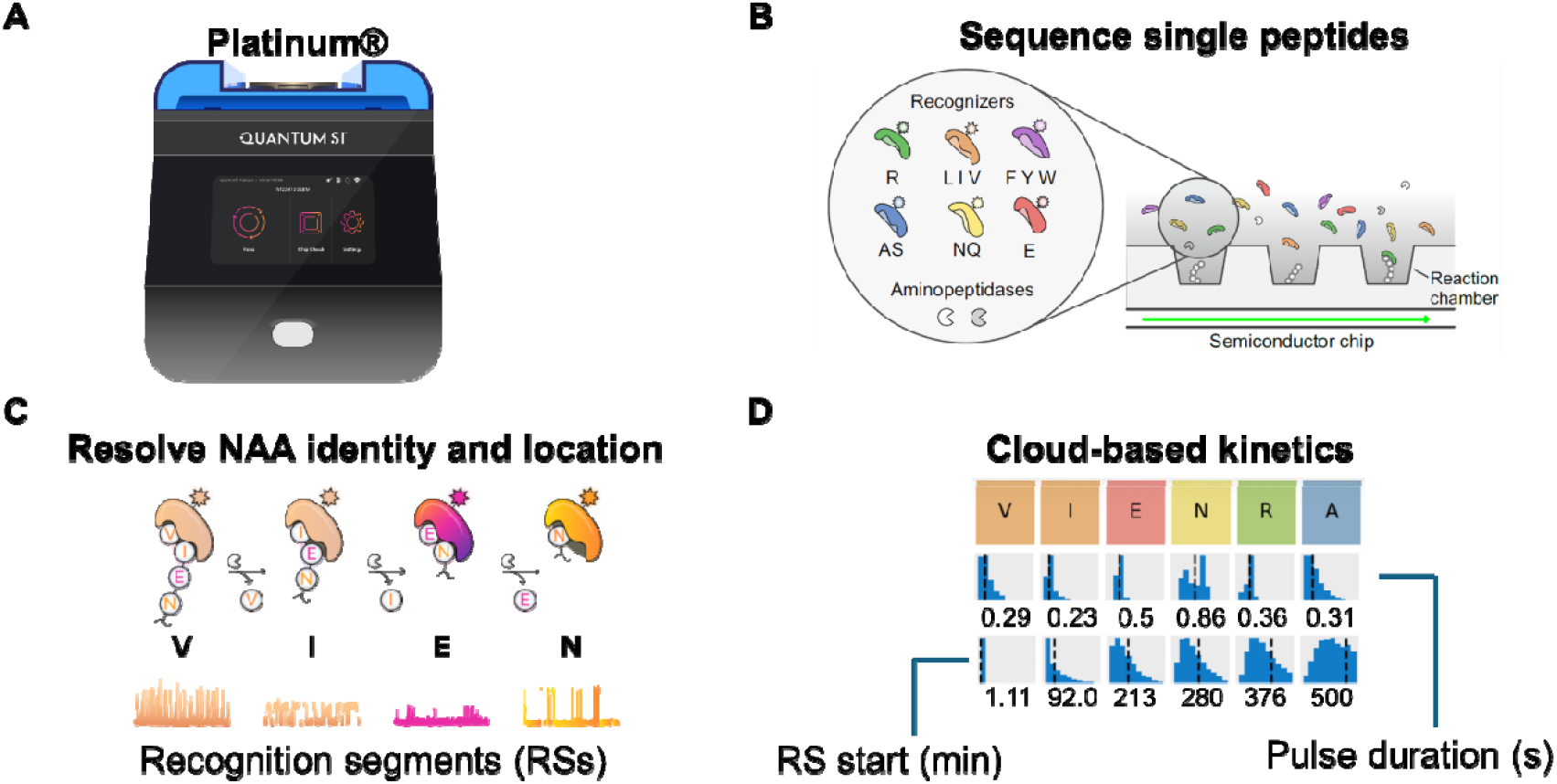
Overview of sequencing on Platinum. (**A**) The Quantum-Si NGPS platform consists of a semiconductor chip, sample prep and sequencing kits, the compact Platinum instrument (27 lbs), and cloud-based software analysis. (**B**) The semiconductor chip contains discrete nanoapertures spread across two flow cells that can accommodate single peptide molecules. The sequencing kits employ aminopeptidases and six N-terminal amino acid (NAA) recognizers, which are labeled with different fluorophores. (**C**) During sequencing, recognizers reversibly bind NAAs, generating characteristic pulsing regions called recognition segments (RSs). As NAA recognizers can target 1-3 NAAs, distinct pulse duration (PD) provides a kinetic readout of NAA identity. (**D**) Upon completion of a sequencing run, data is automatically transmitted for cloud-based analysis. Reads are then aligned to the reference profile based on the correspondence of observed RSs to the expected reference profile, using recognizer identity. Kinetic signature numerical values (RS start, PD) represent the median of a distribution of data across analyzed apertures that contains 4 RS and 3 unique dyes.

### Cloud-based analysis of sequencing data; analysis versions

Primary Analysis v2.5.1 and Peptide Alignment v2.5.2 were used for cloud-based analysis of sequencing data. Details can be found in the Platinum Analysis Software Data Sheet (February 2, 2024). Upon completion of sequencing run, the fluorescent signal, derived from the repeated on-off binding events of the recognizers and NAAs, is securely transferred to the Platinum Analysi Software. The Primary Analysis workflow is the first step in processing data which characterize the apertures across the chip based on peptide loading, recognizer activity, recognizer reads, and recognizer read lengths (**Supplementary Figure 3**). For the Peptide Alignment workflow, reads are aligned to the reference profile based on the correspondence of observed recognition segments to the expected reference profile, using recognizer identity.

## Results and Discussion

We first sought to identify proteoform-specific peptides in TPM1 and 2 to serve as the focus of this study. We performed an in silico digestion of TPM1 and TPM2 with Lys-C, an enzyme that cleaves peptide bonds at the C-terminal side of lysine (K) residues^42^. We then selected eight peptides that correspond to distinct regions along TPM1/2 primary structure (**Figure 1**). In addition, amino acid mutations in these peptides have been found to be relevant to TPM pathophysiology^24–29^ (**Figure 1 and Supplementary Figure 2**).

To discriminate gene paralogs at the peptide level, we tested two sets of paralogous, isobaric peptides on Platinum: ENALDRAEQAEADK (TPM1)/ENAIDRAEQAEADK (TPM2) (**Figure 1 Peptide Pair A, Figure 3A**) and VIESRAQK (TPM1)/VIENRAMK (TPM2) (**Figure 1 Peptide Pair B, Figure 3C**). ENALDRAEQAEADK and ENAIDRAEQAEADK are identical in amino acid sequence except for the 4^th^ position from the N-terminus, in which there is an isobaric Leu/Ile substitution (**Figure 1 Peptide Pair A, Figure 3A**). In other words, they share the same length and molecular weight, and so specifically detecting each peptide requires differentiation of signals from Leu and Ile. We conjugated the peptides and mixed the resulting linker-derivatized peptides in an equimolar ratio for loading on both flow cells of the semiconductor chip (**Methods**). For comparison of ENALDRAEQAEADK and ENAIDRAEQAEADK, the first three amino acids (Glu1/E1, Asn2/N2, Ala3/A3) are identical and each can be measured via distinct recognizers (**Figure 2B**). The recognizer for Leu, Ile, and Val has the highest affinity for N-terminal Leu and this preference is expected to result in markedly longer PD for LDR compared to IDR in these two peptides^21^.

**Figure 3.**
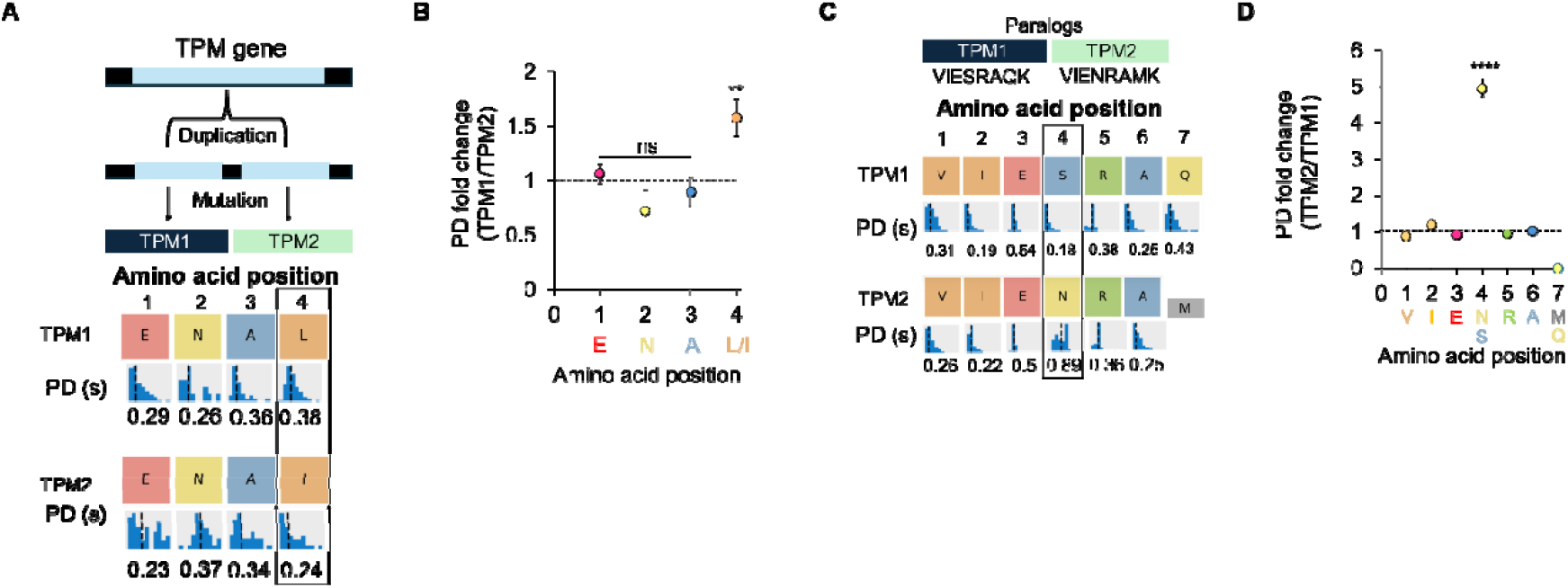
Kinetic signatures discern the identity and amino acid ordering of paralog derived peptides. (**A**) Gene duplication events produce TPM*1*/TPM*2* paralogs, wherein a single isobaric L to I substitution differentiates ENALDRAEQAEADK from ENAIDRAEQAEADK (top). An example kinetic signature snapshot from cloud-based analysis is shown below the amino acid positions (bottom). Colors of the boxes indicate the recognizer identity (**see Figure 1B**). Kinetic signatures such as PD represent the statistical distribution of kinetic data for all pulses associated with a specific residue. (**B**) Plot of the fold change in the average pulse duration (PD) between NAA measurements of ENAL versus ENAI, for the first 4 residues. Error bars indicate standard deviation from three experiments conducted on independent flow cells. Asterisks indicate p-values relative to a PD fold change of 1. (**C)** Gene duplication events produce TPM*1*/TPM*2* paralogs that can be differentiated at position 4 of TPM*1* peptide VIESRAQK and TPM*2* peptide VIENRAMK. (**D**) Plot of the fold change PDs for VIENSRAQ versus VIENRAM. A ∼5-fold higher PD was detected for the N in position 4 of VIENRAMK, relative to S in VIESRAQK, which was statistically significant, when compared to the PD fold change to all other positions. Data (*mean ± S.D.) are representative of three independent experiments, conducted on independent flow cells (ns p >0.05, * p <0.05, ** p <0.01, *** p<0.001, *** p<0.0001; two-tailed t-test). PD=Pulse Duration

Indeed, we observed a higher average PD for Leu (0.35 s) compared to Ile (0.23 s) (**Figure 3B**). These results indicate a single recognizer distinguishes isobaric residues that differ at position 4 from the N-terminus, thus confidently discriminating these paralog-specific peptides.

Interestingly, we observed similar trends of highly specific peptide recognition with the paralogous peptides VIENRAMK and VIESRAQK, in which all residues are the same except for the Asn to Ser substitution in position 4 and the Met to Gln substitution in position 7 (**Figure 1 Peptide Pair B, Figure 3C**). While all matched residues between the two peptides exhibit similar PD profiles, the Asn to Ser substitution at position 4 was distinguished via the dye call for each recognizer (**Figure 3C**). In addition, the recognizers for N and S elicited different PDs for their cognate residues, providing an additional metric for distinguishing each residue at this position. Indeed, a ∼5-fold longer PD was detected for Asn relative to Ser (**Figure 3D**).

In addition to paralogs, the TPM gene family also produces alternatively spliced isoforms that are tissue specific and functionally distinct. In TPM2, exon 6 or 7 is included, but not both, with the same mutually exclusive splicing of exons found for exon 10 and 11. We found that SLMASEEEYSTK maps to a specific splice junction that overlaps exon 6, and thus could inform the presence of the spliceform TPM2-210, which has been found to be expressed in non-skeletal muscle^44–47^ (**Figure 1 Peptide Pair C, Figure 4A**). From the sequencing of SLMASEEEYSTK, PDs were generated for all amino acids within the peptide, except for the Met at position 3 (**Figure 4B**).

**Figure 4.**
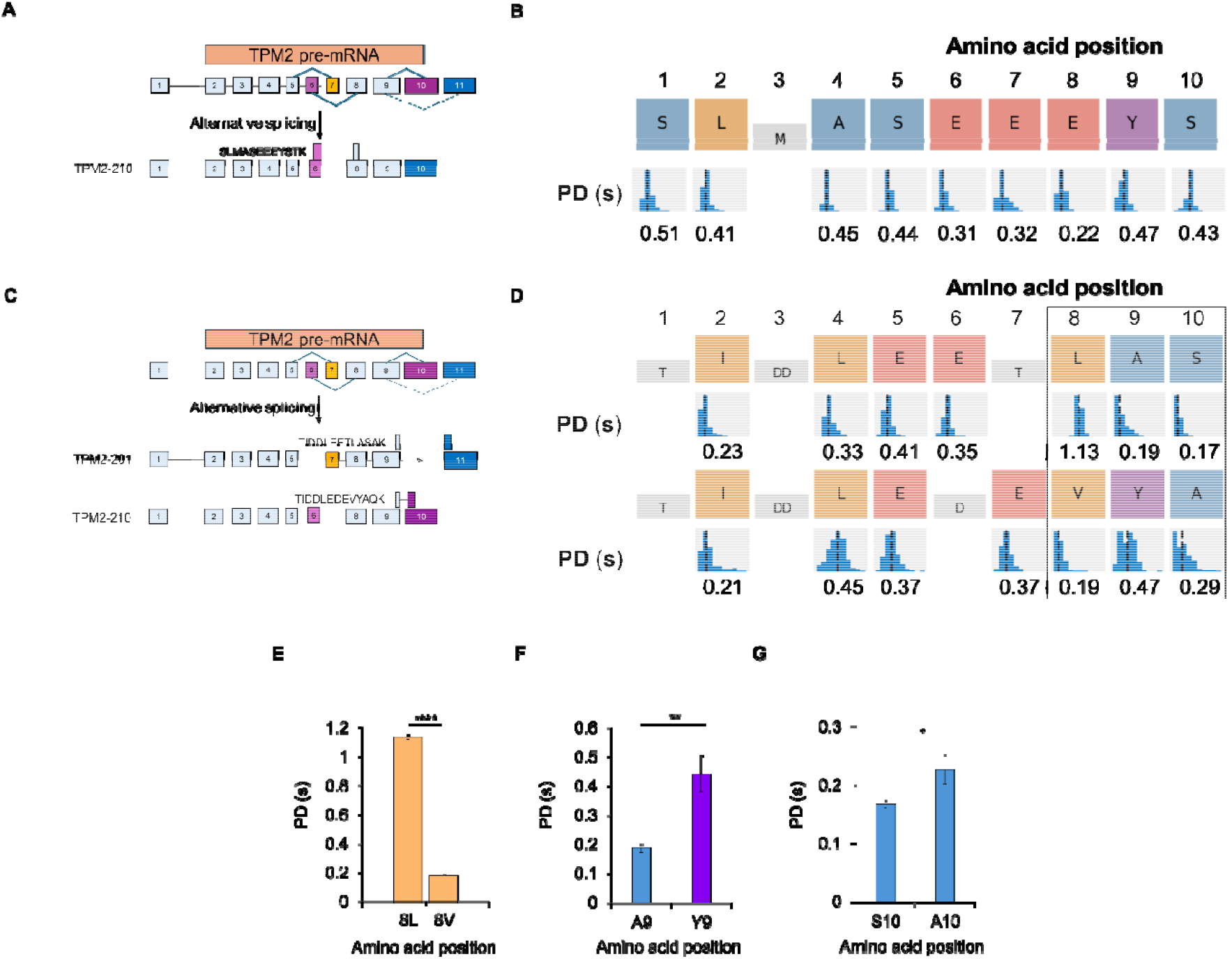
Kinetic signatures discern the identity and temporal order of TPM2 peptides derived from alternative splicing. (**A**) Alternative splicing of TPM2 mRNA generates a tissue-specific spliceform (containing Exon 6) to which SLMASEEEYSTK specifically maps. (**B**) Sequence alignments for SLMASEEEYSTK. An example kinetic signature snapshot from cloud-based analysis is shown below the amino acid positions. Colors of the boxes indicate the recognizer identity (see **Figure 1B**). Kinetic signatures such as PD represent the statistical distribution of kinetic data for all pulses associated with a specific residue. (**C**) Alternative splicing of TPM*2* mRNA differentiates exon-specific peptides TIDDLEETLASAK (Exon 10) and TIDDLEDEVYAQK (Exon 11). (**D**) Sequence alignments for peptides TIDDLEETLASAK and TIDDLEDEVYAQK. (**E**) LIV recognizer elicits differential PD profiles for similar aliphatic residues. A ∼9-fold difference in PD was for observed for L/V. (**F**) Distinct recognizers for A and Y discern position 9. (**G**) The A/S recognizer enables discernment via differential PDs at position 10. Data (*mean ± S.D.) are representative of three independent experiments, conducted on independent flow cells (ns p >0.05, * p <0.05, ** p <0.01, *** p<0.001, *** p<0.0001; two-tailed t-test). PD=Pulse Duration

We also found a peptide pair that discriminates exon 10 from exon 11, with exon-specific peptides of the same length: TIDDLEETLASAK (exon 11) and TIDDLEDEVYAQK (exon 10) (**Figures 1 Peptide Pair D, 4C**). The combination of residue position as well as recognizer identity and PD led to a highly specific discrimination of the peptides (**Figure 4D**). In particular, residues in positions 8-10 were clearly distinguished by recognizer and PD patterns. For position 8—in an additional demonstration of the ability of the LIV recognizer to discern aliphatic amino acids (**Figure 3B**)—we observed a ∼9-fold higher PD for the Leu relative to valine (Val; V) (**Figures 4E**). In position 9, Tyr and Ala residues were clearly distinguished with distinct recognizers (**Figures 4F**). In further support of the ability of a single recognizer to elicit differential PD profiles, we observed a higher PD for Ala relative to Ser in position 10 (**Figures 4G**). Taken together, these results demonstrate specific sequencing of splice junction-specific peptides with Platinum, and that Platinum differentiates peptides based on both structurally similar and distinct amino acid side chains.

Finally, we tested the ability of Platinum to distinguish peptidoforms^48^ arising from PTMs—a major source of proteoform variation. One of the benefits of single-molecule sequencing is discrete binding events between each recognizer and its cognate amino acid, which is a highly sensitive modality of detection. We found that the previously sequenced peptide TIDDLEDEVYAQK (**Figure 4**), which is specific to exon 10 of TPM2, contains an annotated phosphorylation at the Tyr in position 10^49^. The addition of a phosphate group significantly changes the charge and topology of Tyr, weakening binding by the recognizer for bulky aromatic residues (**Figures 1B**).

We separately conjugated TIDDLEDEVYAQK and TIDDLEDEVpYAQK (**Figure 5A**), then loaded each peptidoform for on-chip sequencing in independent flow cells. As expected, we observed a significant decrease in percent of total nanoapertures containing Tyr RSs for the phosphorylated form relative to the unmodified form, while the percent of nanoapertures containing LIV and E RSs were similar between the two groups **(Figure 5B**). These results indicate that Platinum sequencing is sensitive to the presence of PTMs; thus, kinetic signatures can be used to infer the presence of PTMs at singe amino acid resolution.

**Figure 5.**
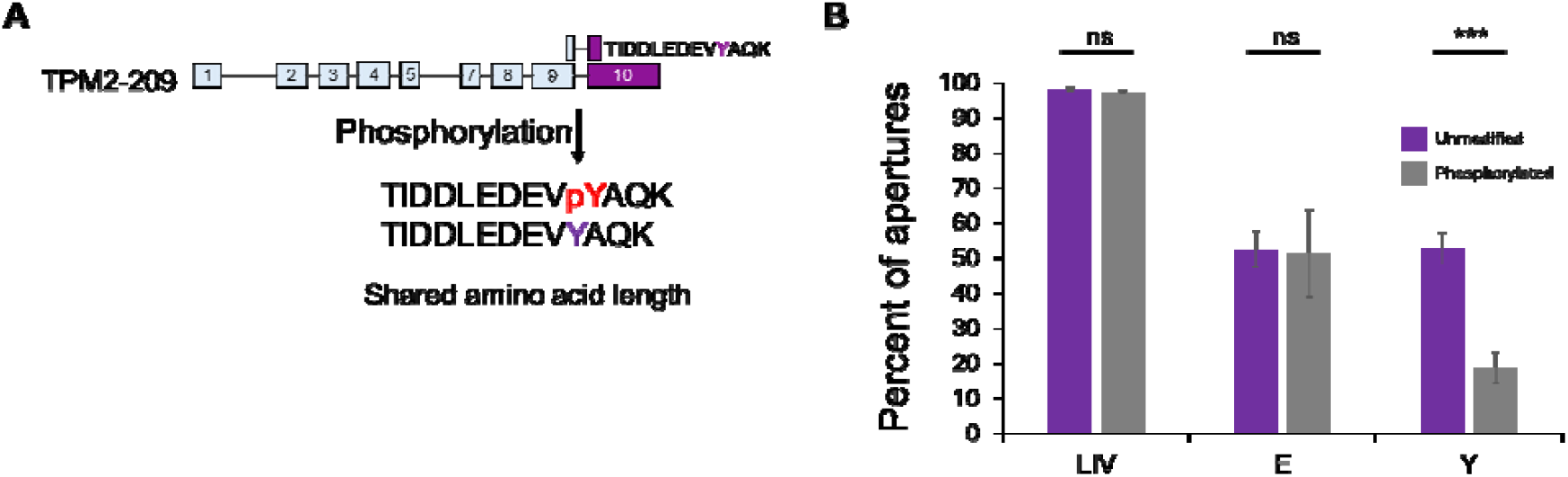
YFW recognizer generates Y-specific recognition events, enabling differentiation from phosphotyrosine-modified peptidoforms. (**A**) Schema showing the exon-specific peptide TIDDLEDEVYAQK, which contains a known phosphosite at the tyrosine residue. (**B**) Bar plot showing the difference in percent of apertures between the recognizers (LIV, E, and FYW) in the unmodified versus phosphorylated version (TIDDLEDEVpYAQK). Data (mean ± S.D.) are representative of three independent experiments, conducted on independent flow cells (ns p >0.05, *** p<0.001; Welch two-sample t-test).

## Conclusions

Analyzing the proteome is essential for a comprehensive understanding of biological processes. However, several fundamental challenges hinder the ability to fully characterize proteomic diversity. These challenges stem from the complexity of the proteome, which could contain over 1 million distinct proteoforms^1,14^.

Existing methods for detecting proteoform-informative peptides, such as mass spectrometry, can struggle with distinguishing closely related variants, particularly those that differ by single amino acid substitutions or PTMs. These methods can have other limitations, including high costs and detection thresholds, as well as the requirement for specific expertise to operate and interpret MS experiments. In contrast, by directly sequencing individual peptides in an accessible benchtop format, Platinum has the potential to overcome some of the limitations of traditional methods. serving as an orthogonal tool for proteomic discovery.

In this study, we sought to determine the ability of the benchtop Platinum instrument to distinguish proteoform-informative peptides derived from the human TPM gene family. Using this single-molecule protein sequencing technology, we identify key variations at the paralog. transcript, and PTM levels. While prior work^21^ laid the foundation for the detection of amino acid variation using Platinum, this study represents the first application of Platinum for sequencing proteoform-informative peptides.

We demonstrate the ability of Platinum to differentiate between paralogous TPM1 and TPM2 peptides that differ by a single amino acid substitution, as well as to sequence tissue-specific TPM2 splice variants. As TPMs are subject to post-transcriptional regulation (e.g. long non-coding RNA and antisense transcripts)^11,50^, there is a need to validate TPM expression at the peptide/protein level. In future studies with biological samples, a pan-TPM antibody that recognizes a conserved site on each TPM isoform could be used for upfront enrichment, followed by Platinum sequencing to identify peptides that uniquely map to each TPM variant. More generally, given its orthogonal chemistry, Platinum can provide additional evidence to support peptide identifications via DDA from MS (**Supplementary Figure 4A and B**).

Our findings also illustrate the sensitivity of Platinum to phosphotyrosine modifications at the single amino acid level. Given this capability, future studies could use Platinum to perform target identification for phospho-specific antibodies^51^. Detecting phosphorylation is particularly challenging due to the transient nature of these modifications and the complex interplay of signaling networks, making high-resolution methods essential for capturing these dynamic processes^52^.

There are several limitations to this study. First, we chose to focus on proteoform-informative synthetic peptides as a model system for this foundational proof-of-concept evaluation. Thus, future studies will be required to extend these findings to biological samples. Second, because we used either 1:1 mixtures of peptides or individual peptides loaded in independent flow cells (Y and pY), future studies with complex mixtures will be required to simulate conditions typically obtained from proteomic studies. Beyond this study, several innovations are needed to expand the biological applications of Platinum, including the development of additional recognizers to expand sequencing coverage. Recently, Quantum-Si released a combined Asp/Glu recognizer; as tropomyosin proteoforms contain an abundance of acidic residues (**Figure 1 and Supplementary Figure 5**), additional peptides could likely be resolved by differentiating acidic sites with the latest sequencing kit.

Broadly, the application of Platinum for proteoform differentiation is generalizable to other proteoform families. While TPMs have been greater than 500 million years in the making^36^, new TPM discoveries are on the horizon, with not only mammals but also six TPM genes that have been found in zebrafish, a popular animal model for gene expression studies^53^. In addition, TPMs are the primary allergens in shellfish, causing food allergies that affect 2% of the US adult population^54,55^. Hence, detecting TPM peptides that elicit food allergies could be of great interest to the food science community^40^.

More generally, NGPS on Platinum could integrate with existing proteogenomic approaches to identify immunopeptides derived from noncoding regions^56,57^, enabling detection of functional biomolecules often overlooked by conventional methods. Further, combining NGPS with genomic and transcriptomic data could provide a more comprehensive view of proteoform diversity, including low-abundance variants and post-translational modifications. This multiomics strategy will lead to an improved understanding of the proteome in health and disease.

## Author contributions

Conception or project design: NS, KAS, EDJ, GMS Data acquisition: NS, KAS, EDJ, GMS

Data analysis and interpretation of results: NS, KAS, EDJ, EFW, GMS Original draft preparation: NS, KAS, EDJ, GMS

All authors reviewed the results and approved the final version of the manuscript.

## Disclosures

KAS is an employee and shareholder of Quantum-Si. No other authors have any disclosures.

## Acknowledgments

We thank members of the Sheynkman Lab, particularly Leon Sheynkman (University of Virginia) for technical assistance. Also, we thank Cheng Man Lun, Meredith Carpenter, and Brian Reed for helpful discussions (Quantum-Si).

## Supplementary data

**Supplementary Figure 1.**
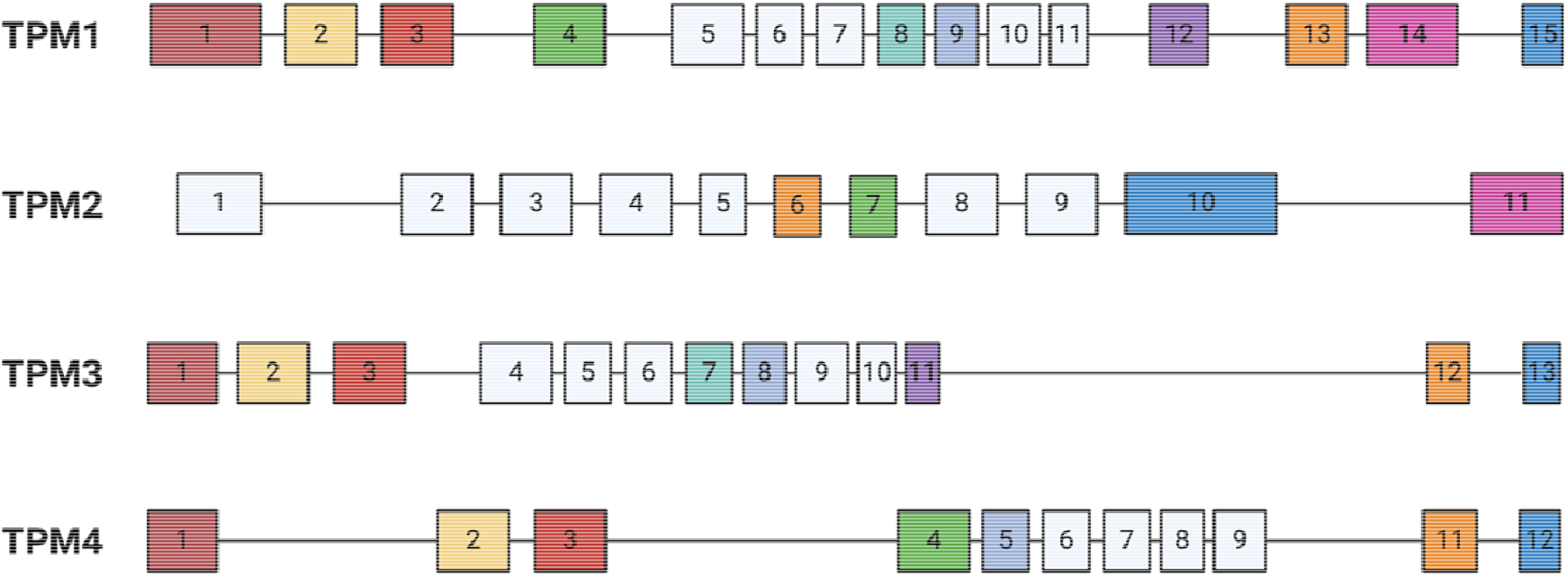
Exonic structure of the human tropomyosin genes. Most vertebrates have four tropomyosin genes, each encoding multiple isoforms due to alternate exon expression. Each tropomyosin gene is composed of 11 to 15 exons, with different splicing patterns correlated with different functions. The *white* boxes are constitutively expressed in all spliceforms of the gene, while *colored* boxes are alternatively spliced to produce distinct spliceforms. The most variable regions of tropomyosin are the ends, encoded by alternate start and end exons that form the intermolecular overlap junction.

**Supplementary Figure 2.**
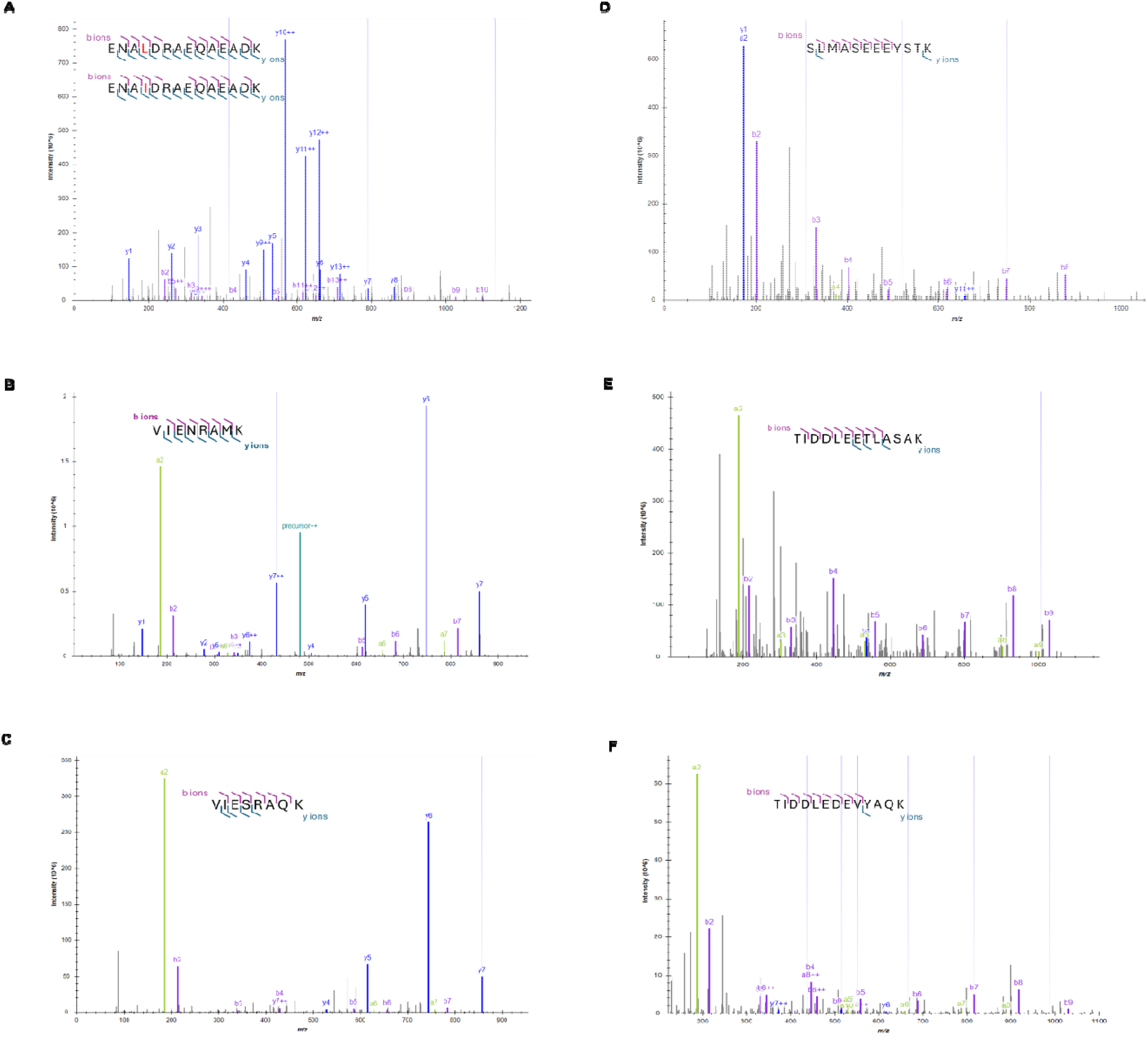
MS2 spectra and fragmentation map of 7 synthetic TPM peptides. **A-F** shows the ion intensities and fragmentation of each peptide: (**A**) ENA[L/I]DRAEQAEDK (spectra are indistinguishable from each other), (**B**) VIENRAMK, (**C**) VIESRAQK, (**D**) SLMASEEEYSTK, (**E**) TIDDLEETLASAK, (**F**) TIDDLEDEVYAQK VIESRAQK, (**D**) SLMASEEEYSTK, (**E**) TIDDLEETLASAK, (**F**) TIDDLEDEVYAQK

**Supplementary Figure 3.**
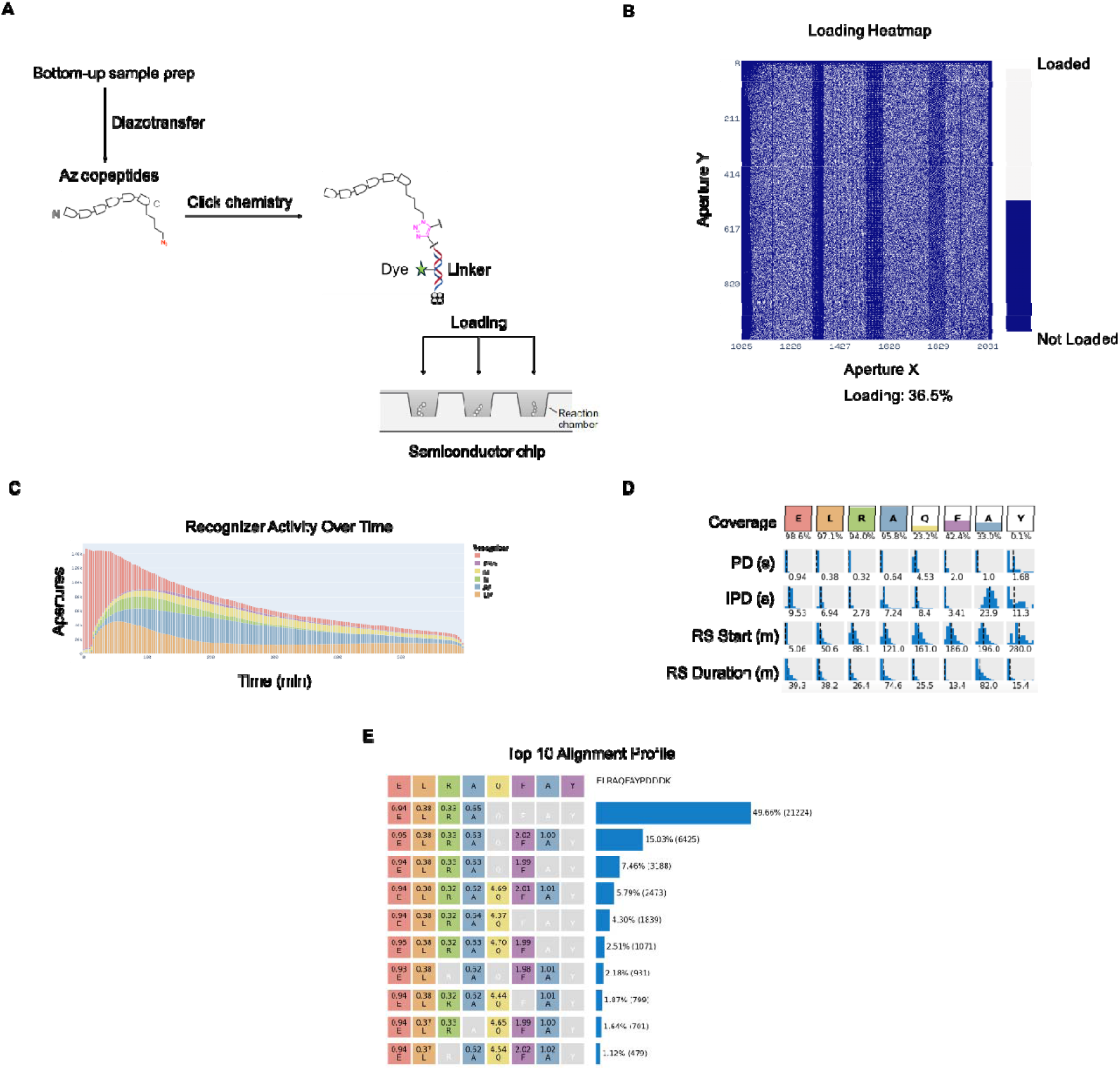
Schematic of Platinum library preparation, chip loading, and peptide alignment. (**A**) Peptides are subjected to diazotransfer to install an N_3_ group at the side chain of the C-terminal lysine. The derivatized peptides are conjugated to fluorescently labeled linkers via click chemistry, then loaded onto flow cells on a semiconductor chip and immobilized in nanoscale reaction chambers (or apertures). (**B**) A heatmap of aperture occupancy is generated after on-chip peptide loading. White pixels represent loaded apertures, while dark blue pixels are unoccupied apertures. (**C**) Recognizer activity over the course of a sequencing experiment. Number of apertures registering pulsing activity for each recognizer type are recorded and plotted in real-time. (**D**) Example coverage map of a synthetic peptide ELRAQFAYPDDK. Colored boxes show the frequency of detection for each residue. The distribution and median of each of the kinetic parameters collected by Platinum are displayed as blue histograms underneath each residue. (**E**) Reads are grouped into different alignment profiles based on the RSs they exhibit. The number of reads in each alignment profile for this peptide are displayed as blue bars to the left of the profiles.

**Supplementary Figure 4.**
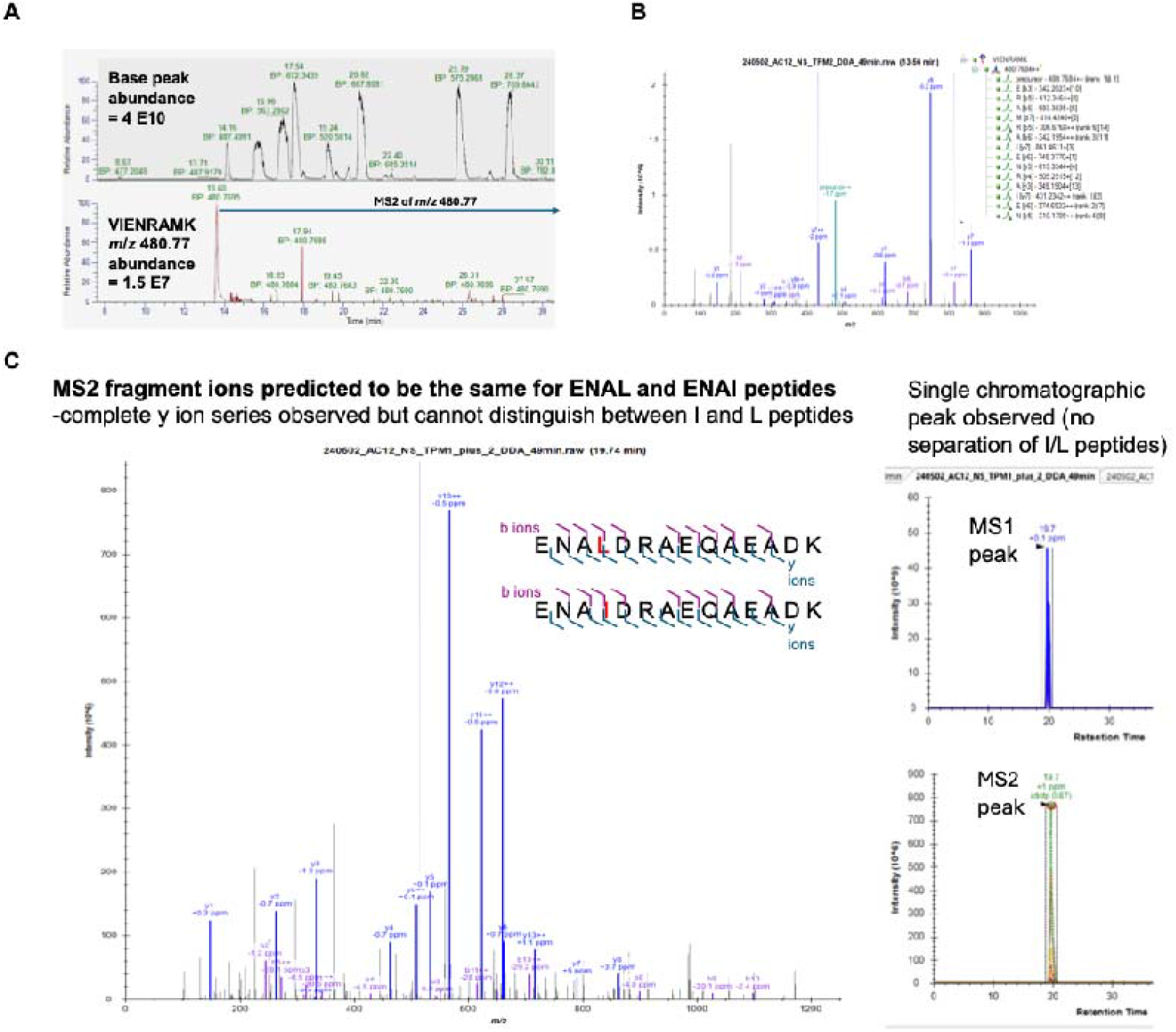
Recombinant TPM2 subjected to endoproteinase Lys-C and LC-MS/MS (Orbitrap Eclipse). (**A**) VIENRAMK peptide ion abundance is 1000-fold less than base peak abundance, leading to a single MS2 scan acquired during DDA method. (**B**) A spectrum of VIENRAMK with b/y ion series. (**C**) Single chromatographic peak observed with isobaric peptides ENALDRAEQAEADK (TPM1) and ENAIDRAEQAEADK (TPM2) complicates LC-MS/MS detection of paralog-specific peptides.

**Supplementary Figure 5.**
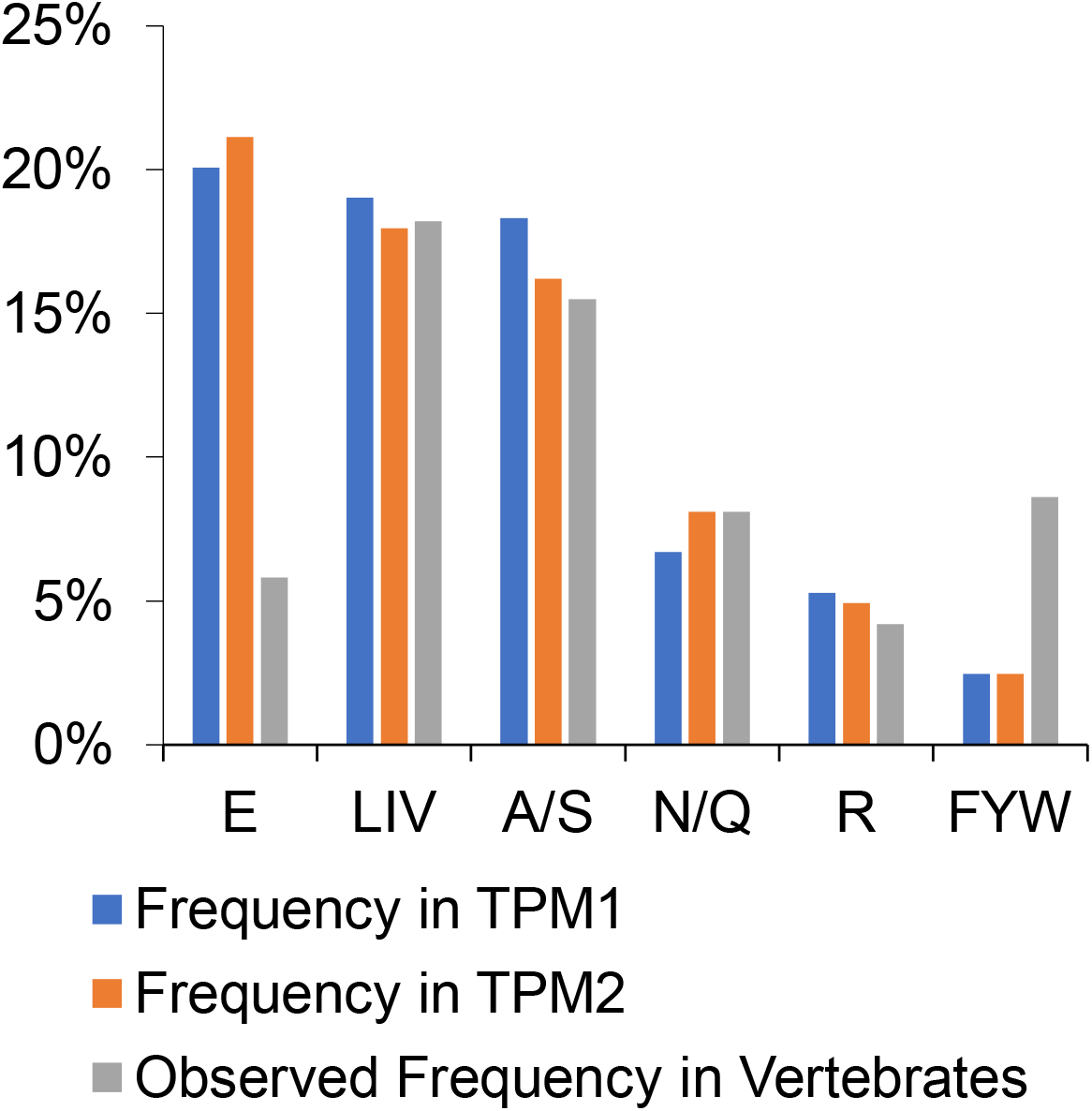
TPM1/2 amino acid frequency of NAA recognizers compatible with Platinum.

